# Cells protect chromosome-microtubule attachments, independent of biorientation, using an Astrin-PP1 and CyclinB-CDK1 feedback loop

**DOI:** 10.1101/2020.12.24.424312

**Authors:** Duccio Conti, Xinhong Song, Roshan L. Shrestha, Dominique Braun, Viji M Draviam

## Abstract

Defects in chromosome-microtubule attachment can cause chromosomal instability, associated with infertility and aggressive cancers. Chromosome-microtubule attachment is mediated by a large macromolecular structure, the kinetochore. Kinetochore pairs are bioriented and pulled by microtubules from opposing spindle poles to ensure the equal segregation of chromosomes. Kinetochore-microtubule attachments lacking opposing-pull are detached by Aurora-B/Ipl1; yet, how mono-oriented attachments that are a prerequisite for biorientation, but lacking opposing-pull are spared is unclear. Using an RNAi-mediated screen, we uncover a unique role for the Astrin-SKAP complex in protecting mono-oriented attachments. We provide the first evidence for how a microtubule-end associated protein senses outer-kinetochore changes specific to end-on attachments and assembles into an outer kinetochore crescent to stabilise mature attachments. We find that Astrin-PP1 and Cyclin-B-CDK1 activities counteract each other to preserve mono-oriented attachments. Thus, cells are not only surveying chromosome-microtubule attachment errors, but they are also actively sensing and stabilising mature attachments independent of biorientation.

---

Non-biooriented kinetochore-microtubule attachments predominate during early mitosis (1–3) and are a step towards biorientation. Biorientation allows the pulling and segregation of chromosomes into two equal sets. Unresolved non-bioriented attachments such as syntelic (co-oriented), monotelic (mono-oriented) and merotelic (multi-oriented) attachments can cause chromosome missegregation. It is known that syntelic and merotelic attachments are resolved through multiple mechanisms: destabilisation of incorrect attachment (Aurora-B/Ipl-1 mediated error correction pathway (4–7)), progressive restriction of attachment geometry (Dynein powered corona-stripping (8–12)) and microtubule-pulling or tension associated active stabilisation of attachments13–15. However, whether and how cells recognise and regulate mono-oriented kinetochore-microtubule attachments is not known.

The significance of monotelic kinetochore geometry and orientation in ensuring normal timing of biorientation and in preventing the formation of syntelic or merotelic attachment errors are evident in several in silico models (16–18). Mono-oriented end-on attachments form in two ways: when kinetochores interacting with microtubule-walls become tethered to microtubule-ends (end-on conversion (19–21)), and when microtubules nucleate from kinetochores as pre-formed K-fibers (22,23). Oscillations of mono-oriented chromosomes (including velocity, duration and amplitude) are similar to other chromosome oscillations (2,3,24), but how mono-oriented attachments are protected without opposing-pull is unclear. One potential way to protect a mono-oriented kinetochore-microtubule attachment from Aurora-B mediated error-correction could be by establishing tension through its sister kinetochore’s interaction with microtubule-walls, as lateral kinetochore-microtubule interactions are immune to Aurora-B action (25,26). However, this is unlikely to be the primary mechanism as stable monotelic kinetochores lacking lateral microtubule interaction have been observed in a variety of conditions (21,23,24). Other kinetochore-bound kinases, CDK1 and PLK1, have been implicated in stabilising chromosome-microtubule attachments (27) but their impact specifically on mono-oriented attachments (independent of microtubule-mediated opposing pull) remain unclear.

Here we present evidence for how human cells exploit a localised kinase-phosphatase feedback loop to preserve mono-oriented end-on attachments independent of biorientation. Through a targeted RNAi-screen, we uncovered the role of the Astrin-SKAP complex in preserving mono-oriented attachments even when the error-correction pathway is switched off. To understand how Astrin facilitates the sensing or stabilisation of mono-oriented attachments, we used time-lapse microscopy tools to assess Astrin arrival and departure at kinetochores. We found the role of CDK1 in regulating Astrin enrichment at end-on attachments, which is important for the ‘sensing’ but not the ‘error-correction’ of attachments. To explain how Astrin senses end-on attachments, we performed deletion studies and identified a short 273a.a region of Astrin sufficient to recognise outer-kinetochore changes selective to end-on attached kinetochores. We show that the Astrin-delivered pool of PP1 dephosphorylates a subset of CDK1 phosphorylation sites on Bub1 but not CENP-T or Ndc80, revealing a mechanism linking the arrival of Astrin and departure of Mad1-Mad2. Thus Astrin-PP1 and CyclinB-CDK1 form a negative feedback loop to maintain non-bioriented attachments, separate from the canonical Aurora-B mediated pathway for error correction.

## Results

### Astrin-SKAP protect mono-oriented attachments independent of Aurora-B pathway

We find that blocking the recruitment of BubR1-B56 phosphatase, an enzyme crucial for end-on attachments in bipolar spindles (25,28), disrupts the maintenance of mono-oriented end-on attachments (Supplementary Fig. 1a - 1c). This shows that phospho-signalling is likely to underpin the life of mono-oriented kinetochore-microtubule attachments. Hence, we set out to explore the how Aurora-B mediated error-correction of syntelic and merotelic attachments coexist with the need to preserve monotelic attachments. we performed an RNAi-based screen of kinetochore proteins, particularly those delivering phosphatases, to identify those required to maintain non-bioriented end-on attachments even in the absence of Aurora-B. We depleted kinetochore-microtubule bridging proteins using standardised siRNA oligos (25,29–31) (Supplementary. Table 1) and exposed cells to an Eg5 inhibitor (STLC or Monastrol) to induce monopolar spindles followed by a brief exposure to ZM443479 and MG132 to inhibit Aurora-B and block anaphase onset, respectively. In control cells, this treatment results in monopolar spindles with kinetochores tethered to the tip of microtubule-ends forming a bouquet-like arrangement (Supplementary Fig. 1d).

**Figure 1.**
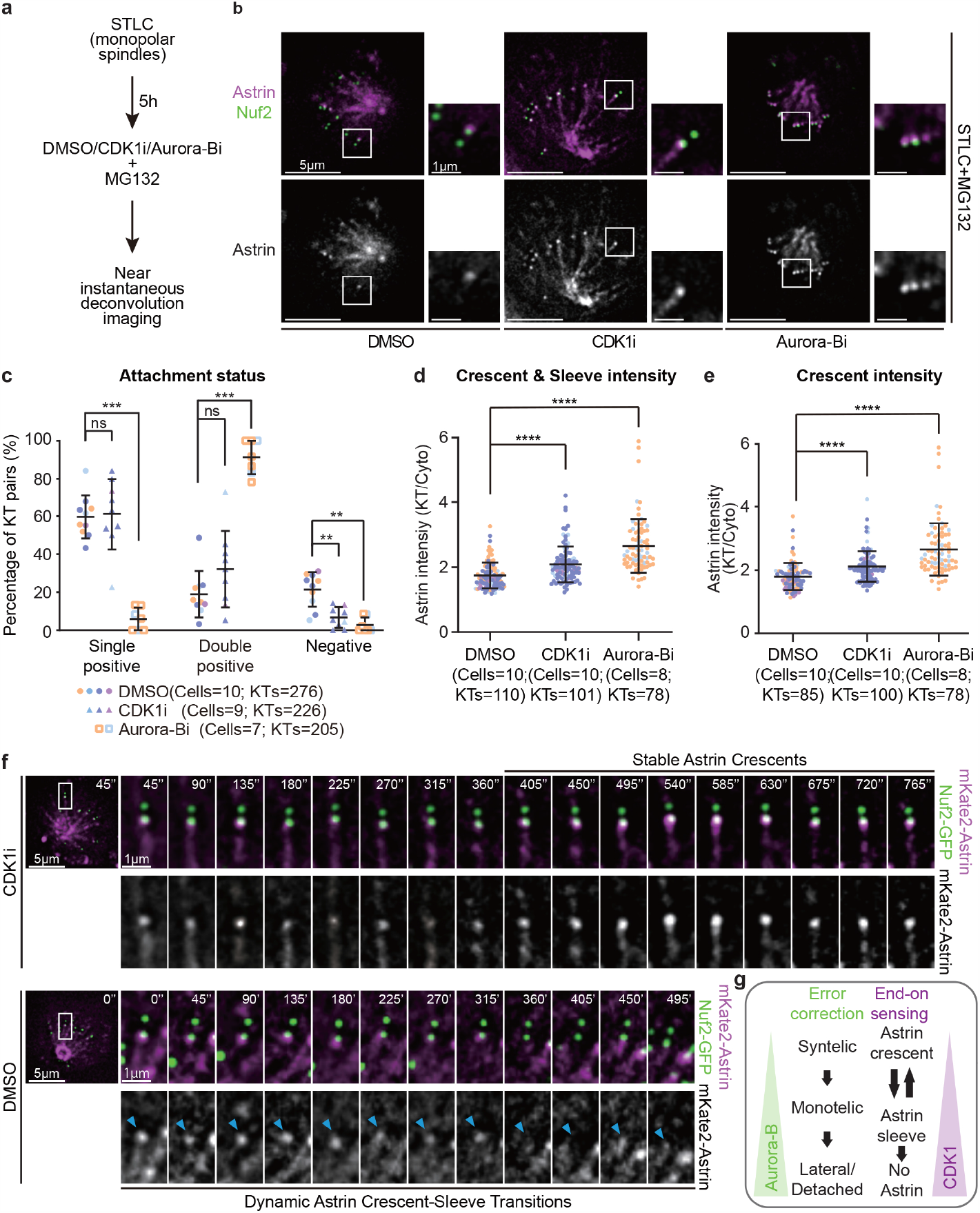
CDK1 activity controls Astrin-mediated sensing of end-on attachment. a, Experimental regime shows near-instantaneous live-imaging of kinetochore fate soon after the addition of kinase inhibitors to understand end-on attachment sensing mechanisms in non-bioriented kinetochores. b, Live cell images of monopolar spindles show monotelic kinetochores soon after DMSO or RO3666 treatment and syntelic kinetochores soon after ZM447439 treatment in cells co-expressing Nuf2-GFP and mKate2-Astrin. Cells were exposed to STLC for 5 hours before adding CDK1 (RO3306) or AuroraB (ZM447439) inhibitor or DMSO (control) with the proteasome inhibitor, MG132. Scale bar as indicated. c, Graph shows the percentage of monotelic (Astrin single-positive) or syntelic (Astrin double-positive) kinetochore pairs following inhibitor treatment as indicated using images as shown in b. Colours in super-plot represent independent experimental repeats. d and e, Graph shows intensities of either Astrin crescents and sleeves (d) or Astrin crescents alone (e) in cells treated as in a. Colours in super-plot represent independent experimental repeats. f, Time lapse images of kinetochores show changes in Astrin intensities at kinetochores of monopolar spindles in cells coexpressing mKate2-Astrin and Nuf2-CFP and treated with DMSO but not CDK1 inhibition. Cells were treated with STLC for 5 hours and MG132 added along with either CDK1 inhibitor or DMSO (control) as indicated and filming started soon after drug addition. Time interval between frames is 45 seconds. Scale as indicated. g, Illustration of separable roles for Aurora-B and CDK1 in regulating Astrin-mediated sensing of end-on attachments. CDK1 decreases the amount of Astrin at kinetochores without disrupting error-correction that is dependent on Aurora-B.

We assessed attachment status using two methods: (i) comparing kinetochore position at the end of microtubule (end-on interaction) resulting in a monopolar bouquet-like arrangement of kinetochores and (ii) the presence or absence of Astrin crescents, a marker specific for stable end-on kinetochores independent of biorientation (25,32). In this targeted screen, codepletion of Ndc80-Nuf2 was used as a positive control since the Ndc80 complex is essential for all forms of kinetochore-microtubule attachments. Depletion of chTOG1 (33,34), Astrin-SKAP complex, SKA complex, EB1-EB3 complex and CLIP-170 - all known to bind both the kinetochore and microtubule-ends (reviewed in 27), resulted in aberrant nuclei (multinucleated or misshapen) as expected, confirming chromosome missegregation induced by protein depletion (Supplementary Table 1). Our analysis of attachment status showed that chTOG1, EB1-EB3 complex, SKA complex and CLIP170 are not essential for maintaining end-on attachments in the absence of Aurora-B activity (Supplementary Table 1 and Supplementary Figure 1d). Uniquely Astrin or SKAP depleted cells showed a reduction in the number of end-on attached kinetochores and loss of bouquet-like chromosome arrangement in monopolar spindles in this screen (Supplementary Fig. 1d; Supplementary Table-1). Thus, the RNAi-screen showed Astrin-SKAP as a Kinetochore-associated MAP needed to maintain non-bioriented end-on attachments even in the absence of Aurora-B.

### CDK1 activity controls Astrin-mediated sensing of end-on attachment

The heterotetrameric Astrin-SKAP complex enriches specifically at end-on (and not lateral) kinetochores (21). We hypothesised that factors that control Astrin recruitment at non-bioriented kinetochores can reveal how cells sense and stabilise end-on attachments before biorientation. So, we searched for factors that control Astrin enrichment by studying the incidence and intensity of Astrin at kinetochores of monopolar spindles in fixed-cell studies (Supplementary fig. 2a and 2b). As expected inhibiting the error-correction enzyme Aurora-B increased the percentage of Astrin double-positive kinetochore pairs showing an increase in syntelic attachments (Supplementary Fig. 2c). In addition, Aurora-B inhibition increased the amount of Astrin at kinetochores (Supplementary Fig. 2d), as reported (35). Surprisingly, CDK1 inhibition increased the amount of Astrin at the outer-kinetochore (Supplementary Fig. 2d), without promoting syntelic attachments (Supplementary Fig. 2c). Thus CDK1 activity influences Astrin-mediated sensing of end-on attachments without disrupting error correction. To confirm these findings, we analysed live-cells soon after the addition of CDK1 inhibitor

**Figure 2.**
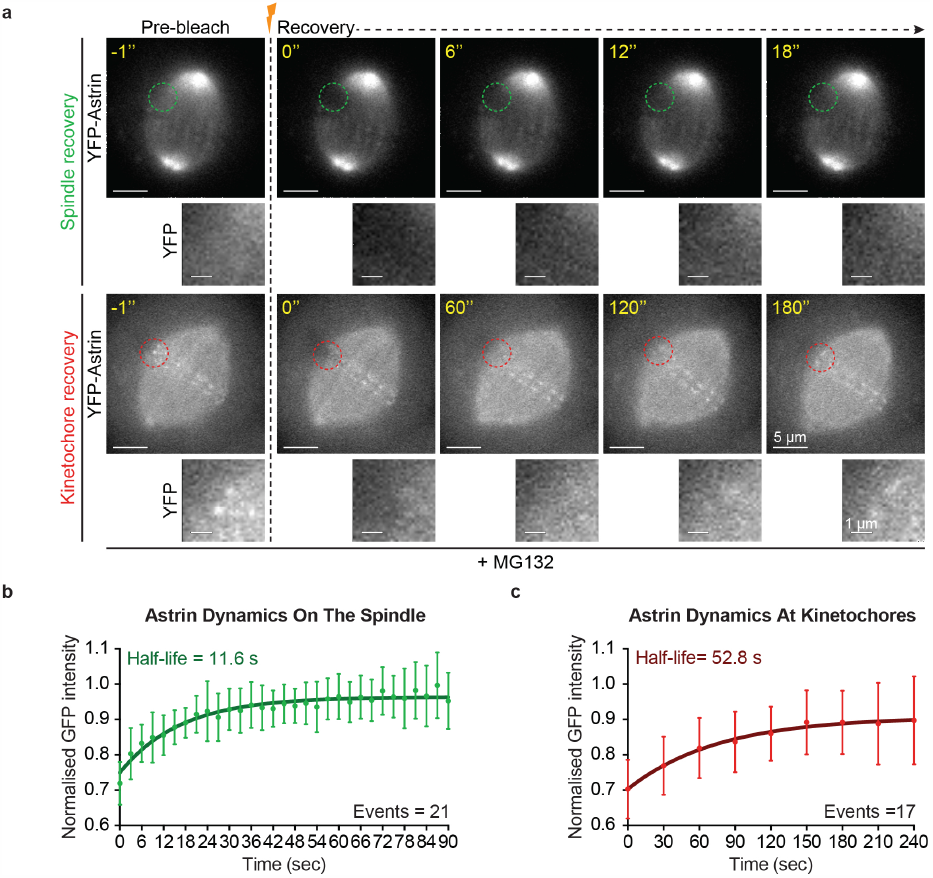
Autonomous sensor of end-on attachment at individual kinetochores. a, Representative time-lapse images of Fluorescence Recovery After Photobleaching (FRAP) of HeLa cells expressing YFP-Astrin following one hour of MG132 treatment prior to photobleaching. Cropped images show the recovery of YFP intensities after bleaching of one spindle area or one kinetochore highlighted with green or red dotted circles, respectively. Scale bar as indicated. b and c, Graphs show curves of normalised YFP fluorescence intensities on spindle (b) or kinetochores (c), respectively, versus time from FRAP experiments as in a. Lines and whiskers mark the average and standard deviation, respectively, from two independent experiments (kinetochore data) and three independent experiments (microtubules data). The calculated halflife time of recovery is indicated in each graph. The turnover rate of kinetochore-bound Astrin is at least 4-fold higher compared to turnover rates of GFP-tubulin at kinetochore bound microtubules (7-9 min (37–39)), suggesting that the dynamic outer kinetochore shell of Astrin is not directly related to microtubule plus-end assembly rates. Consistent with this idea, the turnover rate of YFP-Astrin on spindle microtubules is somewhat similar to the turnover rate of GFP-tubulin in non-kinetochore microtubules (21.5 sec (37)). Thus, the distinct pool of Astrin at kinetochores, and the rapid turnover of Astrin at kinetochores compared to the turnover of tubulin at kinetochore-microtubules show that Astrin forms a dynamic outer-kinetochore crescent when chromosomes are end-on attached.

(Fig. 1a). In monopolar spindles of cells co-expressing mKate2-Astrin and Nuf2-GFP (a kinetochore marker) and exposed to DMSO (solvent control), 80% of kinetochores pairs displayed mKate2-Astrin (Fig. 1b and 1c), of which 80% were monotelic (single-positive) and 20% were syntelic (double-positive) (Fig. 1c). Consistent with fixed-cell studies (Supplementary Fig. 2), the amount of kinetochore-bound Astrin and the percentage of syntelic attachments were increased following Aurora-B inhibition (Fig. 1c and 1d). In contrast, CDK1 inhibition increased the amount of kinetochore-bound Astrin (Fig. 1d) but did not significantly increase the percentage of syntelic attachments (Fig. 1c), confirming CDK1’s role in controlling Astrin recruitment without disrupting error-correction. We have shown that Astrin localisation fluctuates between a crescent and sleeve shape at the outer-kinetochore of monopolar spindles32. We quantified level changes in Astrin crescents alone and observed a significant increase in Astrin crescent intensities following CDK1 inhibitor treatment (Fig. 1e). Thus, both fixed and live-cell studies show that unlike Aurora-B, CDK1 activity regulates Astrin enrichment at end-on kinetochores without interfering with error correction mechanisms. CDK1 is reported to be required for establishing kinetochore-microtubule attachments (36). So, we were surprised to observe an increase in the amount of kinetochore-bound Astrin following CDK1 inhibition, which suggests a role for CDK1 in blocking Astrin-mediated sensing of mono-oriented end-on kinetochores (Fig. 1f). Therefore, to investigate how CDK1 activity influences Astrin enrichment at end-on kinetochores, we tracked mono-oriented kinetochores soon after the addition of CDK1 inhibitor using deconvolved time-lapse microscopy of monopolar spindles of cells coexpressing mKate2-Astrin and Nuf2-YFP. Careful temporal analysis of change in Astrin levels at mono-oriented end-on kinetochores showed dynamic changes in Astrin signals (crescent versus sleeve) in control (DMSO-treated) cells, as reported in unperturbed cells32 (Fig. 1f). However, in CDK1 inhibitor treated cells, kinetochore crescents of Astrin did not transition into sleeves, within 5 minutes of CDK1 inhibitor treatment (Fig. 1f). Thus, Astrin enrichment at mono-oriented end-on attachment is counteracted by CDK1. In summary, (i) Astrin-mediated dynamic sensing of non-bioriented end-on kinetochores is under the influence of CDK1, (ii) CDK1 inhibition induced Astrin enrichment does not impair error-correction and (iii) CDK1 and Aurora-B kinase activities influence the fate of kinetochore-microtubule attachments differently (Fig. 1g).

### Autonomously regulated outer-kinetochore pool of Astrin crescents

Unlike mono-oriented end-on kinetochores where Astrin crescents and sleeves alternate dynamically, bioriented end-on kinetochores stably display Astrin crescents (Fig. 1 and (32)). To explore whether Astrin-mediated sensing of end-on attachments continues to be dynamic at bioriented kinetochores, we measured Astrin turnover. Fluorescence Recovery After Photobleaching (FRAP) of YFP-Astrin showed that the recovery of YFP-Astrin signal intensities at kinetochore was significantly slower compared to spindle microtubules: 52.8 s (range of 29.5 - 252.4 s) at kinetochores versus 11.6 s (range of 9.6 - 14.6 s) on microtubules (Fig. 2a-2c). This indicates two distinct pools of Astrin: one at the outer kinetochore and the other along spindle microtubules. Importantly, the FRAP rates reveal a wide range in the half-lives of YFP-Astrin at kinetochores, compared to microtubules, indicating autonomous changes in Astrin turnover at individual kinetochores.

### A 273 a.a region of Astrin senses outer-kinetochore changes specific to end-on attachments

How cells sense end-on attachments independent of biorientation is not clear, although it is known that Mad2 and Mad1 leave the kinetochore soon after the formation of end-on attachment (20,21). Astrin-SKAP is enriched at end-on but not lateral kinetochores (25), making it a strong candidate for directly recognising outer kinetochore changes specific to end-on attachments, and so we searched for the minimal kinetochore targeting domain in Astrin. The C-terminal tail of Astrin is needed to deliver PP1 and not localise at kinetochores32. To identify the region of Astrin needed to recognise outer-kinetochore changes specific to end-on attachments we exploited GBP-PP1 (GFP-Binding Protein fused to PP1) to constitutively deliver PP1 near Astrin-GFP’s C-termini(32) (Fig. 3a). We first asked whether the N-terminus of Astrin (1-693 a.a.) that forms the Astrin-SKAP tetramer and binds to microtubule-walls (40) is essential for the complex’s kinetochore localisation. Surprisingly, neither the Astrin-SKAP interaction domain nor Astrin’s microtubule-binding domain is needed to target Astrin at metaphase kinetochores, as 694-1122 a.a of Astrin-GFP can be recruited to kinetochores when coexpressed with GBP-PP1 (Fig. 3b). Notably, this recruitment required GBP-PP1 and could not be achieved simply by blocking Aurora-B activity (data not shown). Through further deletions, we narrowed down a 273 a.a stretch (851-1122 a.a) of Astrin as the shortest kinetochore targeting region sufficient for recognising structural changes at the outer-kinetochore specific to end-on attachments. Interestingly, both the 694-1122 and 851-1122 a.a fragments of Astrin, co-expressed with GBP-PP1, showed a reduction in the incidence and intensities of crescents compared to Astrin full-length protein (Fig. 3c and Supplementary Fig. 3).

**Figure 3.**
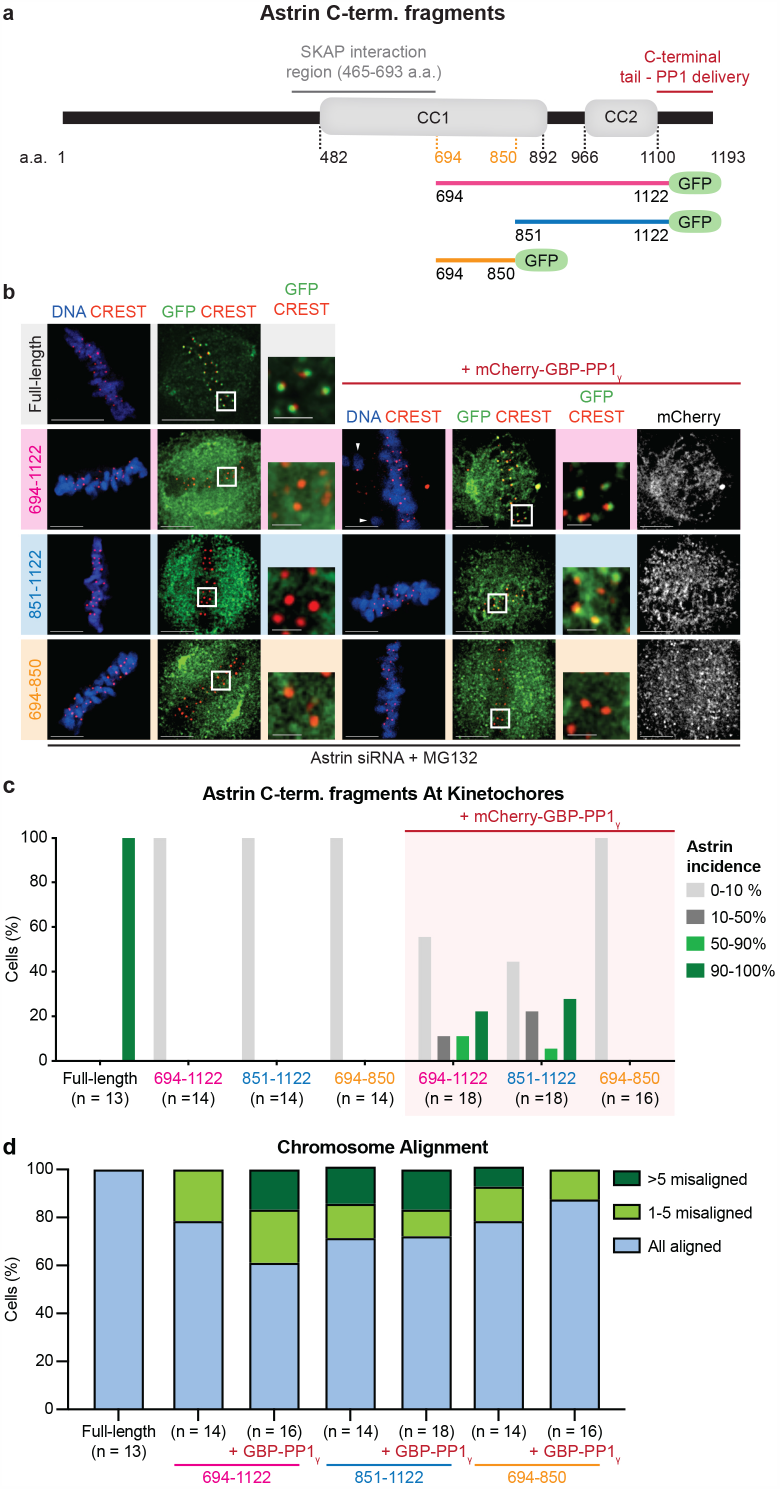
A 273 a.a region of Astrin senses outer-kinetochore changes specific to end-on attachments. a, Schematic of Astrin fragments used to ascertain minimal kinetochore targeting domain of Astrin. b, Images of immunostained cells show the localisation of GFP tagged Astrin fragments (as indicated) in the presence or absence of mCherry-GBP-PP1 following an hour of MG132 treatment prior to fixation. Cells were immunostained with antibodies against GFP, mCherry and CREST (a kinetochore marker). White boxes mark the area of cropped images. Scale 5 and 1 microns in main and cropped images, respectively. White arrowheads mark unaligned chromosomes quantified in Supplementary Fig. 3. c, Bar graphs show the percentage of cells displaying GFP-tagged fragments of Astrin, as indicated, in the form of outer-kinetochore crescents at 90-100% 50-90% 10-50% or 0-10% of aligned kinetochores. Cells co-expressing GBP-PP1 are marked separately. d, Bar graphs show the percentage of cells with chromosome alignment defects in cells expressing GFP-tagged fragments of Astrin with or without mCherry-GBP-PP1 and arrested in MG132 for an hour before immunostaining with antibodies against GFP, mCherry and CREST (a kinetochore marker). To assess the extent of misalignment, cells were segregated into three bins: all chromosomes aligned, 1-5 misaligned chromosomes and more than 5 misaligned chromosomes.

These findings shows that although 851-1122 a.a. is sufficient to recognise outer kinetochore changes, the microtubule-binding N-terminus of Astrin is important for the complete outer-kinetochore crescent formed by the Astrin complex.

Metaphase cells expressing Astrin fragments (694-1122 or 851-1122 a.a) showed several unaligned kinetochores, highlighting the need for the full outer-kinetochore shell of Astrin to ensure stable maintenance of congressed chromosomes (Fig. 3d). In summary, to sense outer-kinetochore changes specific to end-on attachments, the microtubule-interacting region of Astrin is dispensable, although this region works closely with the end-on kinetochore sensing region to build full outer-kinetochore crescents of Astrin at mature end-on kinetochores.

### Astrin delivered PP1 lowers Bub1 phosphorylation linked to Mad1 departure

To understand how mono-oriented attachments are protected from the premature action of error-correction pathways, we searched for downstream substrates of Astrin-PP1 at kinetochores. A signal transduction cascade, including MPS1-KNL1(MELpT)-Bub1-Mad proteins, operates closely attenuating checkpoint signals induced by MPS1 (reviewed in 41). While we know how this check-point silencing cascade responds to the departure of checkpoint kinases (41), we do not know how this cascade responds to the arrival of Astrin-PP1 phosphatase at end-on kinetochores.

The feedback loop involved in rapidly silencing the checkpoint is likely to be closely tied to end-on attachment sensing mechanisms, as Mad2 and Mad1 checkpoint proteins are known to rapidly leave end-on kinetochores before biorientation (20,21). So, we took advantage of Astrin-4A mutant with a crippled PP1-docking site (32) as a molecular probe to disrupt Astrin-mediated PP1 delivery and study its impact on phospho-events at the outer-kinetochore. Immunostaining for phospho-epitopes showed that cells expressing Astrin-WT or Astrin-4A mutant display no significant changes in the incidence or intensity of HEC1-pSer55 (Aurora-B substrate)42 or CENP-T-pSer47 (CDK1 substrate)43 at kinetochores (Fig. 4a and Supplementary Fig. 4a-4b and 4g-4h). However, compared to Astrin-WT expressing cells, Astrin-4A mutant expressing cells display a noticeable increase in KNL1(MELpT) (44,45) and Bub1(pSpT)(46) positive kinetochores (Supplementary Fig. 4c and 4e and Fig. 4a).

**Figure 4.**
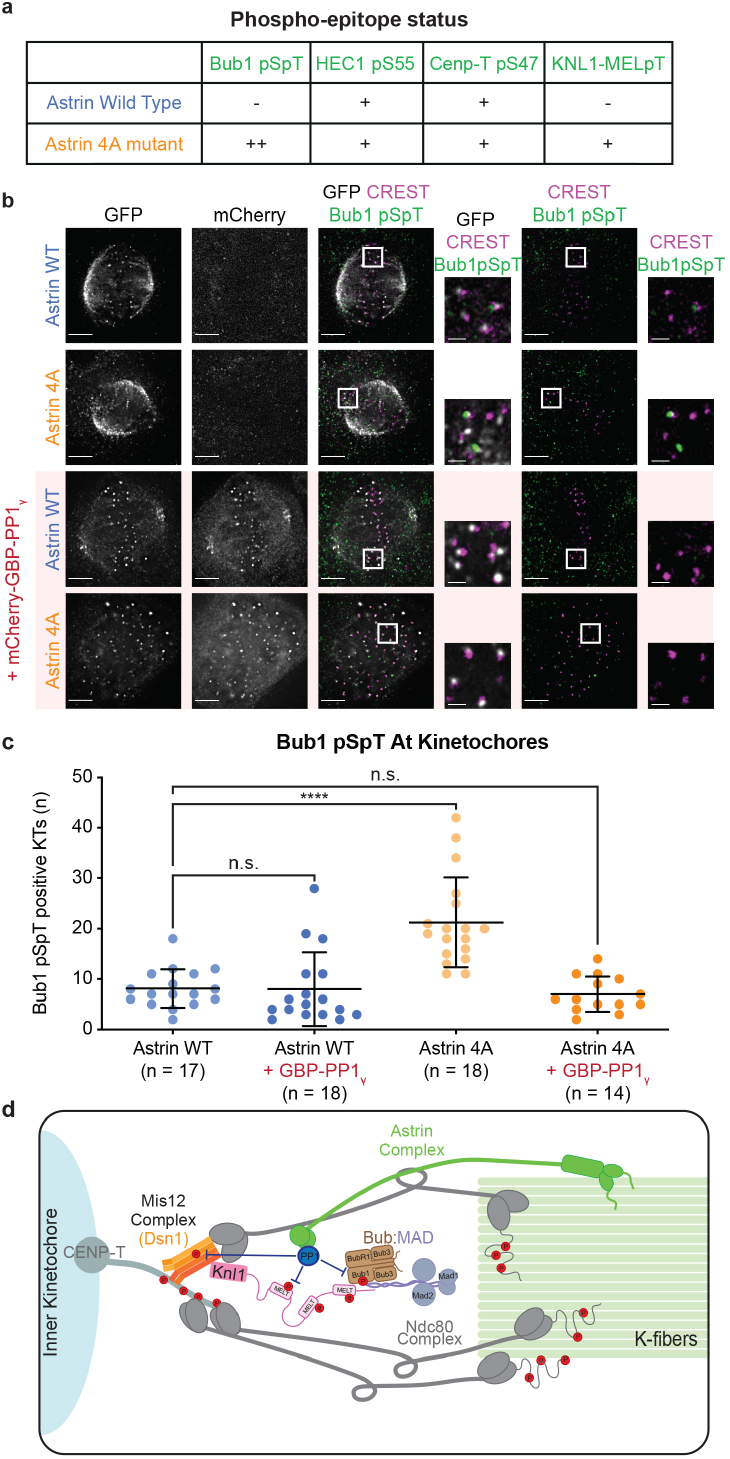
Astrin delivered PP1 lowers Bub1 phosphorylation linked to checkpoint silencing. a, Table summarises the phosphorylation status of various phospho-epitopes at the outer kinetochore. ‘+’ or ‘-’ refers to phospho-epitope observed in more or less than 20% of kinetochores, respectively, as shown in Supplementary Fig. 4. b, Images show the extent of phosphorylated Bub1 (pSpT) at kinetochores of cells depleted of endogenous Astrin and expressing either Astrin-WT or Astrin-4A mutant alone or along with mCherry-GBP-PP1 as indicated. Cells were treated with MG132 for an hour prior to fixation for immunostaining with antibodies against GFP and phospho-epitopes as indicated and CREST antisera (a centromere marker). Scale bar. 5 microns. c, Graph of the number of kinetochores positive for phosphorylated Bub1 (pSpT) determined using whole spindle Z-stacks of images as shown in b. Each dot represents values from one cell. Bars and whiskers represent mean and standard deviation, respectively, across data from three experimental repeats. “*” and “ns” indicate significant and insignificant statistical differences, respectively, as calculated using a non-parametric Kruskal-Wallis H test combined with Dunn’s multiple comparisons test.d, Model of dephosphorylation events that depend on the recruitment of Astrin-PP1 at end-on kinetochores. Astrin-PP1 alters the phosphorylation of (i) Bub1 (brown) responsible for positioning Mad1 near KNL1-MELT, (ii) DSN1 of Mis12 complex (orange) and (iii) KNL1-MELpT (pink) sites but not those of Ndc80 (S44) or CenpT (S47) sites (in grey), indicating a spatially restricted dephosphorylation of substrates.

The increase in the proportion of Bub(pSpT) and KNL1(MELpT) positive kinetochores (Supplementary Fig. 4d and 4f) is in agreement with the increase in Mad2 and Zw10 displaying metaphase kinetochores in Astrin-4A expressing cells (32). So, we tested whether the increase in levels of phosphorylated Bub1 (pSpT; CDK1 substrate (46)) in Astrin-4A-GFP expressing cells could be rescued by coexpressing GBP-PP1. Immunostaining studies show a reduction in Bub1(pSpT) positive kinetochores in cells coexpressing Astrin-4A-GFP and GBP-PP1 compared to those expressing Astrin-4A-GFP alone (Fig. 4b and 4c), confirming a rescue of Astrin-4A mutant phenotype. Thus, a pool of PP1 phosphatase delivered by Astrin can dephosphorylate a subset of outer-kinetochore substrates and thereby control the fate of mono-oriented kinetochores by working closely with the KNL1-Bub1-MAD1 signalling cascade (Fig. 4d).

### CDK1 and Astrin-PP1 counteract each other at the outer-kinetochore

The dephosphorylation of Bub1 by Astrin-PP1 (Fig. 4) and the enrichment of Astrin following CDK1 inhibition (Fig.1) indicate an opposing relationship between the activities of CDK1 kinase and Astrin-PP1 phosphatase. To test this hypothesis, we first probed whether the arrival of Astrin/SKAP-PP1 and departure of Cyclin-B-CDK1 at kinetochores are correlated. Immunostaining studies of RPE1 cells using antibodies against SKAP and Cyclin-B revealed prometaphase kinetochores displaying either SKAP or Cyclin-B signals (Fig. 5a and 5b), suggesting a departure of Cyclin-B in Astrin-SKAP enriched kinetochores. This was confirmed by quantification of Cyclin-B status at kinetochores enriched for SKAP: Cyclin-B high kinetochores were rarely associated with SKAP-high kinetochores but frequently associated with SKAP-low or SKAP lacking kinetochores (Fig. 5c), demonstrating that CyclinB is significantly reduced at kinetochores enriched for Astrin-SKAP complex.

**Figure 5.**
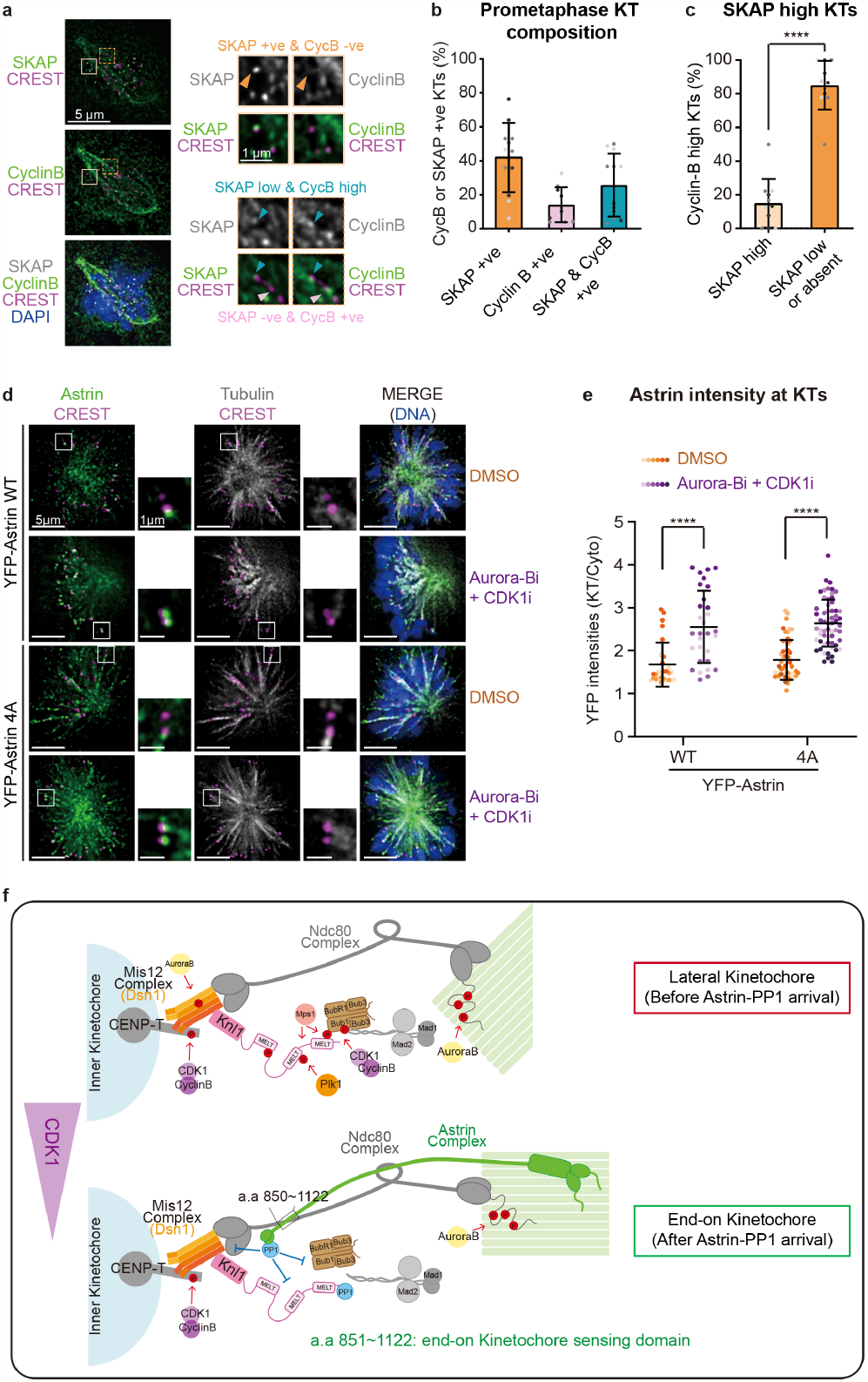
CDK1 and Astrin-PP1 counteract each other at the outer-kinetochore. **a**, Images of RPE1 cells immunostained with antibodies against CyclinB and SKAP and CREST anti-sera and co-stained with DAPI for DNA. Arrowheads mark a kinetochore with low SKAP and high CyclinB intensities (blue), SKAP-positive and CyclinB-negative kinetochore (orange), SKAP-negative and CyclinB-positive kinetochore (pink). Scale as indicated. b, Graph of percentage of kinetochores enriched for either CyclinB or SKAP or both, ascertained using images as in a. c, Graph of percentage of CyclinB positive kinetochores with SKAP-high or -low intensities as in a. Height of bars and whiskers (b and c) mark average value and standard deviation, respectively, across experimental repeats. Each dot represents a value from one cell. d, Images show Astrin-4A mutant at kinetochores of monopolar spindles exposed to Aurora-B and CDK1 inhibitors. Cells depleted of Astrin and expressing GFP-tagged Astrin-4A mutant or -WT as indicated were treated with STLC followed by MG132 and either DMSO or ZM447439 and RO3306 for 30 minutes. Cells were immunostained with antibodies against GFP and Tubulin and CREST anti-sera. Scale as indicated. e, Graph of YFP-Astrin WT or 4A mutant intensities at kinetochores of cells as in d. Black bars and whiskers mark average value and standard deviation, respectively, across experimental repeats. Each dot represents a value from one kinetochore. ‘*’ and ‘ns’ indicate statistically significant and insignificant differences of the Non-parametric Mann-Whitney test, respectively. Colours of dots in b, c, and e represent experimental repeats. f, At end-on kinetochores, Astrin delivers a pool of PP1 that counteracts a subset of spatially-limited phosphorylation events by multiple kinases (CDK1, Aurora-B or PLK1). Inhibiting CDK1 and Aurora-B activities supersedes the need for Astrin-mediated PP1 delivery, revealing a feedback loop that protects non-bioriented end-on attachments from kinase activities.

To test whether Astrin-PP1 and Cyclin-B-CDK1 constitute a negative feedback loop, we checked Astrin-4A localisation in anaphase when CDK1 activity is reduced through CyclinB degradation (50). Immunostaining showed Astrin-4A mutant enrichment at kinetochores in anaphase (Supplementary Fig. 5) but not metaphase (35). This shows that in the absence of CDK1 activity, Astrin-mediated delivery of PP1 is dispensable for Astrin enrichment at end-on kinetochores. To directly test whether one of the main roles of Astrin delivered PP1 is to counteract CDK1 substrate phosphorylation at kinetochore and thereby, retain Astrin at mono-oriented kinetochores, we probed whether CDK1 inhibition will allow the enrichment of Astrin-4A mutant at mono-oriented end-on kinetochores (35). Immunostaining studies showed that Astrin-4A mutant can be maintained at non-bioriented end-on kinetochores following the co-inhibition of CDK1 and Aurora-B activities (Fig. 5d and 5e) but not Aurora-B inhibition alone (35). These show that reducing CDK1 activity can compensate for the loss of Astrin delivered PP1. We conclude that a key role of Astrin delivered PP1 is to counteract CDK1 activity at the outer-kinetochore (Fig. 5f), which in turn promotes Astrin’s enrichment and stabilises non-bioriented kinetochore-microtubule attachments.

## Discussion

Here, we report that non-bioriented end-on attachments are sensed and protected, independent of biorientation mechanisms. We show that Astrin-mediated sensing of end-on attachment is dynamic, and identify a short 200a.a region of Astrin as sufficient to recognise outer-kinetochore changes specific to mature end-on attachments. Of the kinetochore associated MAPs known to deliver PP1, Astrin’s role is unique: different from the role of CENP-E that binds PP1 and delivers CLASP at naked kineto-chores47, Astrin-PP1 can sense mature end-on attachments (Fig. 5f). Although the microtubule-associated SKA complex can enhance Ndc80-microtubule coupling and stability of bioriented attachments48 like Astrin-SKAP complex, neither the SKA complex nor CENP-E is important for maintaining end-on attachment in the absence of Aurora-B (this study). Thus, Astrin’s kinetochore-microtubule interaction (through microtubule-binding N-terminus), its PP1 delivery role (through C-terminal tail) and outer-kinetochore change sensing role (through C-terminal coiled-coil region) bring in a set of unique kinetochore-microtubule bridging function which ultimately stabilise kinetochore-microtubule attachments independent of biorientation.

We show that Astrin enrichment at non-bioriented kinetochores is regulated by CDK1, without compromising error correction mechanisms. Conversely, Astrin-delivered PP1 regulates a subset of CDK1 phospho-sites at the kinetochore, and the lack of Astrin-delivered PP1 can be overcome by counteracting CDK1. These show the entangling of the activities of CDK1 kinase and Astrin-PP1 phosphatase in a feedback loop appropriate for rapidly sensing and stabilising end-on attachments, without interfering with biorientation mechanisms. This is consistent with a role for CDK1 in establishing attachments (reviewed in 27) and Astrin-SKAP complex’s role in maintaining, but not forming, end-on attachments21.

The close association between the mechanism to sense non-bioriented end-on attachments - through Astrin-PP1 enrichment - and the signalling cascade that silences the checkpoint is relevant for checkpoint silencing in meiosis I where homologs are separated while sister kinetochores remain mono-oriented and end-on tethered following end-on conversion49,50. Uncovering spatially localised mechanisms for controlling kinetochore-microtubule attachment stability, without invoking whole-cell changes, is important since chromosomal instability can be restored by tweaking attachment stability that’s lost in most aneuploid cancer cells (39).

## METHODS

### Cell Culture and Drug treatments

HeLa cells (ATCC) cultured in Dulbecco’s Modified Eagle’s Media and RPE1 cells (ATCC) cultured in Ham’s-F12 were supplemented with 10% FCS and antibiotics (Penicillin and Streptomycin). For live-cell imaging studies, cells were seeded onto 4-well cover glass chambered dishes (Lab-tek; 1064716). HeLa and RPE1 cell lines were tested and confirmed free of Mycoplasma. For inhibition studies, cells were treated with 10µM RO3306 (1305, TOCRIS), 20 µM STLC (83265,TOCRIS), 10µM ZM447439 (2458, TOCRIS), 100 nM Taxol (T7191, SIGMA-ALDRICH) or 10µM MG132 (1748, TOCRIS). For monopolar spindle studies, STLC or Monastrol treatment was for 1 to 2 hours and then supplemented with MG132 and kinase inhibitors for 15’ to 30’ prior to fixation. For bipolar spindle studies, MG132 treatment was for 1 hour.

### Plasmid and siRNA Transfection

siRNA transfection was performed using Oligofectamine according to manufacturer’s instructions. Two siRNA oligos were used to target Astrin mRNA 5’ UTR (GACUUGGUCUGAGACGUGAtt) or Astrin 52 oligo (UCCCGACAACUCACAGAGAAAUU). Negative control siRNA (12,935–300) was from Invitrogen. Plasmid transfection was performed using TurboFect (Fisher; R0531) or DharmaFECT duo (Dharmacom; T-2010) according to manufacturer’s instructions. In addition to the standard protocol, after 4 h of incubation, the transfection medium was removed and a fresh selected pre-warmed medium was added to each well. In co-transfection studies, eukaryotic expression vectors encoding Astrin and mCherry-GBP were used in 3:1 ratio. mCherry-GBP-PP1 expression plasmid was generated by subcloning 7-300 of PP1 into an mCherry-GBP expression plasmid. Astrin mutants are described in 35. Plasmid sequences were confirmed by DNA sequencing. Plasmids and plasmid maps are deposited in Ximbio.com.

### Immunofluorescence studies

Cells were cultured on ø13 mm round coverslips (VWR; 631-0150). Unless specified, cells were fixed with ice-cold methanol for a minute. For cold stable assays, 4% PFA was used as in the Super-Resolution Microscopy fixation protocol 28. Following fixation, two quick washes with a wash buffer (1X PBS + 0.1% Tween 20) were performed, followed by three washes of 5 minutes each. Coverslips were incubated with (1X PBS + 0.1% Tween 20 + 1% BSA) for 20 minutes, before staining with primary antibodies overnight at 4°C. For assessing phospho-epitopes, cells were treated with prewarmed buffers. Following a quick rinse with PHEM (60 mM PIPES, 25 mM HEPES, 10 mM EGTA, 4 mM MgSO4, pH 6.9), cells were exposed to fixation buffer (4% PFA in PHEM), and then lysed at 37°C for 5 minutes in PHEM buffer, 1% Triton X-100 and Protease/Phosphatase Inhibitor Cocktail (Cell Signalling Technology; 5872S) before re-exposing to fixation buffer for 20 minutes. All subsequent washes were performed thrice, with 5-minute incubations, in PHEM-T (PHEM buffer + 0.1% Triton X-100) at room temperature. Fixed cells were washed and then blocked with 10% BSA in PHEM buffer for an hour at room temperature before overnight incubation at 4°C with primary antibodies diluted in PHEM buffer with 5% BSA, which was followed by a wash before incubation with secondary antibodies diluted in PHEM + 5% BSA for 45 minutes at room temperature. Finally, coverslips were washed with PHEM-T except before mounting onto glass slides when coverslips were quickly rinsed in distilled water. Cells were stained with antibodies against -Tubulin (Abcam; ab6160; 1:800 or 1:500), Cyclin B1 (Abcam; ab72; 1:1000), GFP (Roche; 1181446001; 1:800), mCherry (Thermo Scientific; M11217; 1:2000), SKAP (Atlas; HPA042027; 1:1000), Astrin (Novus; NB100-74638; 1:1000), GFP (Abcam; ab290; 1:1000), mCherry (Abcam; ab167453; 1:2000), CENP-T pSer47 (Sigma; abe1846; 1:1000), HEC1 pSer55 (Fisher; PA5-85846; 1:500), KNL1 MELpT (Kops Lab; 1:1000), Bub1 pSpT (Nilsson Lab; 1:1000) and CREST antisera (Europa; FZ90C-CS1058; 1:2000) were used. DAPI (Sigma) was used to co-stain DNA. All antibody dilutions were prepared using the blocking buffer. Images of immunostained cells were acquired using 100X NA 1.4 objective on a DeltaVision Core microscope equipped with CoolSnap HQ Camera (Photometrics). Volume rendering (SoftWorx) was performed for 3D analysis of kinetochore-microtubule attachment status (as in 28). Deconvolution of fixed-cell images and 3D volume rendering were performed using SoftWorx*™*.

### Microscopy and Image analysis

For all high-resolution live-cell imaging assays, cells were either transfected with plasmid vectors 24h before imaging or directly transferred to imaging in Leibovitz’s L15 medium (Invitrogen; 11415064) supplemented with MG132 (1748, TOCRIS; 10 µM) and incubated for 1h at 37°C before imaging (as in 53,54 and 55). For high-resolution live-imaging at least 6 Z-planes, 0.3m apart, were acquired using a 100X NA 1.40 oil immersion objective on an Applied Precision DeltaVision Core microscope equipped with a Cascade2 camera under EM mode. Imaging was performed at 37°C using a full-stage incubation chamber set up to allow normal mitosis progression 55 and microtubule dynamics 56. For HeLa cells transiently expressing mKate2-Astrin and Nuf2-CFP, kinetochores were identified by Nuf2-CFP signal. Images were analysed using FIJI Software (NCBI)using a 6×6 pixel circle area. Ratios were calculated using ExcelTM (Microsoft) and graphs plotted using Prism6*™*(*GraphPad*).

### Astrin-PP1 constitutive binding assay

HeLa cells were treated with Astrin siRNA(28), incubated for 24 hours and then transfected with plasmid vectors encoding Astrin-GFP alone or co-transfected with mCherry-GBP-PP1. 24 hours post-transfection of plasmids, cells were arrested for 4 hours in media containing 20 µM STLC, which was followed by a transient exposure to 10 µM ZM447439 and RO3306 and 10 µM MG132 for 15 minutes. Cells were quickly washed thrice with warm DMEM, then washed 5 times with 5 minutes pause using warm DMEM. Cells were fixed 45 minutes after release and immunostained using antibodies against GFP (for Astrin-GFP), mCherry (for mCherry-GBP-PP1) and antibodies against phospho-epitopes and DAPI stain (for DNA).

## Author’s contributions

DC designed and performed experiments in Figures 2, 3, 4 and Supplementary Figures 1b-c, 3, 4 and 5. XS designed and performed experiments in Figure 1, contributed to image acquisition and analysis in Figures 4d and 5 and Supplementary Figures 2 and 4, and generated cartoons in Figures 4 and 5. VMD contributed to three of the experiments within the RNAi screen in Supplementary Table 1. The RNAi screen was designed, conducted and analysed by RLS. Experiments for Supplementary Figure-2 were conducted by DB with the support of RLS. VMD conceived the experiments and wrote the manuscript. XS and DC supported text revision and figure preparation.

## Acknowledgements

We would like to acknowledge funding support from BBSRC (R01003X/1), MRC (MR/K50127X/1), QMUL (SBC8DRA2), Chinese Scholarship Council (CSC file no. 201906820034) and CRUK (C28598/A9787). We would like to thank the Nilsson and Kops laboratories for sharing reagents. We acknowledge Sam Court for infrastructure maintenance support, Christoforos Efstathiou and Parveen Gul for detailed comments on the manuscript, and Draviam group members for discussions on data analysis.

## References

1. Roos, U.-P. Light and electron microscopy of rat kangaroo cells in mitosis. Chromosoma 54, 363–385 (1976).

2. Cassimeris, L., Rieder, C. L. Salmon, E. D. Microtubule assembly and kinetochore directional instability in vertebrate monopolar spindles: implications for the mechanism of chromosome congression. J. Cell Sci. 107 (Pt 1), 285–297 (1994).

3. Bajer, A. S. Functional autonomy of monopolar spindle and evidence for oscillatory movement in mitosis. J. Cell Biol. 93, 33–48 (1982).

4. Biggins, S. Murray, A. W. The budding yeast protein kinase Ipl1/Aurora allows the absence of tension to activate the spindle checkpoint. Genes Dev. 15, 3118–3129 (2001).

5. Tanaka, T. U. et al. Evidence that the Ipl1-Sli15 (Aurora kinase-INCENP) complex promotes chromosome bi-orientation by altering kinetochore-spindle pole connections. Cell 108, 317–329 (2002).

6. Hauf, S. et al. The small molecule Hesperadin reveals a role for Aurora B in correcting kinetochore–microtubule attachment and in maintaining the spindle assembly checkpoint. J. Cell Biol. 161, 281–294 (2003).

7. Ditchfield, C. et al. Aurora B couples chromosome alignment with anaphase by targeting BubR1, Mad2, and Cenp-E to kinetochores. J. Cell Biol. 161, 267–280 (2003).

8. Hoffman, D. B., Pearson, C. G., Yen, T. J., Howell, B. J. Salmon, E. D. Microtubule-dependent changes in assembly of microtubule motor proteins and mitotic spindle checkpoint proteins at PtK1 kinetochores. Mol. Biol. Cell 12, 1995–2009 (2001).

9. Magidson, V. et al. Adaptive changes in the kinetochore architecture facilitate proper spindle assembly. Nat. Cell Biol. 17, 1134–1144 (2015).

10. Sacristan, C. et al. Dynamic kinetochore size regulation promotes microtubule capture and chromosome biorientation in mitosis. Nat. Cell Biol. 20, 800–810 (2018).

11. Wynne, D. J. Funabiki, H. Kinetochore function is controlled by a phospho-dependent coexpansion of inner and outer components. J. Cell Biol. 210, 899–916 (2015).

12. Auckland, P., Clarke, N. I., Royle, S. J. McAinsh, A. D. Congressing kinetochores progressively load Ska complexes to prevent force-dependent detachment. J. Cell Biol. 216, 1623–1639 (2017).

13. Akiyoshi, B. et al. Tension directly stabilizes reconstituted kinetochore-microtubule attachments. Nature vol. 468 576–579 (2010).

14. Dumont, S., Salmon, E. D. Mitchison, T. J. Deformations within moving kinetochores reveal different sites of active and passive force generation. Science 337, 355–358 (2012).

15. Skibbens, R. V., Skeen, V. P. Salmon, E. D. Directional instability of kinetochore motility during chromosome congression and segregation in mitotic newt lung cells: a push-pull mechanism. J. Cell Biol. 122, 859–875 (1993).

16. Mogilner, A. Craig, E. Towards a quantitative understanding of mitotic spindle assembly and mechanics. J. Cell Sci. 123, 3435–3445 (2010).

17. Gay, G., Courtheoux, T., Reyes, C., Tournier, S. Gachet, Y. A stochastic model of kinetochore-microtubule attachment accurately describes fission yeast chromosome segregation. J. Cell Biol. 196, 757–774 (2012).

18. Edelmaier, C. et al. Mechanisms of chromosome biorientation and bipolar spindle assembly analyzed by computational modeling. Elife 9, (2020).

19. Sikirzhytski, V. et al. Microtubules assemble near most kinetochores during early prometaphase in human cells. J. Cell Biol. 217, 2647–2659 (2018).

20. Magidson, V. et al. The spatial arrangement of chromosomes during prometaphase facilitates spindle assembly. Cell 146, 555–567 (2011).

21. Shrestha, R. L. Draviam, V. M. Lateral to end-on conversion of chromosome-microtubule attachment requires kinesins CENP-E and MCAK. Curr. Biol. 23, 1514–1526 (2013).

22. Banerjee, B., Kestner, C. A. Stukenberg, P. T. EB1 enables spindle microtubules to regulate centromeric recruitment of Aurora B. J. Cell Biol. 204, 947–963 (2014).

23. Khodjakov, A., Copenagle, L., Gordon, M. B., Compton, D. A. Kapoor, T. M. Minus-end capture of preformed kinetochore fibers contributes to spindle morphogenesis. J. Cell Biol. 160, 671–683 (2003).

24. Rieder, C. L., Davison, E. A., Jensen, L. C., Cas-simeris, L. Salmon, E. D. Oscillatory movements of monooriented chromosomes and their position relative to the spindle pole result from the ejection properties of the aster and half-spindle. J. Cell Biol. 103, 581–591 (1986).

25. Shrestha, R. L. et al. Aurora-B kinase pathway controls the lateral to end-on conversion of kinetochore-microtubule attachments in human cells. Nat. Commun. 8, 150 (2017).

26. Kalantzaki, M. et al. Kinetochore–microtubule error correction is driven by differentially regulated interaction modes. Nat. Cell Biol. 17, 421–433 (2015).

27. Conti, D. et al. How are Dynamic Microtubules Stably Tethered to Human Chromosomes? Cytoskeleton - Structure, Dynamics, Function and Disease (2017) doi:10.5772/intechopen.68321.

28. Kruse, T. et al. Direct binding between BubR1 and B56–PP2A phosphatase complexes regulate mitotic progression. J. Cell Sci. 126, 1086–1092 (2013).

29. Draviam, V. M., Shapiro, I., Aldridge, B. Sorger, P. K. Misorientation and reduced stretching of aligned sister kinetochores promote chromosome missegregation in EB1-or APC-depleted cells. EMBO J. 25, 2814–2827 (2006).

30. Tamura, N. et al. A proteomic study of mitotic phase-specific interactors of EB1 reveals a role for SXIP-mediated protein interactions in anaphase onset. Biol. Open 4, 155–169 (2015).

31. Grimaldi, A. D. et al. CLASPs are required for proper microtubule localization of end-binding proteins. Dev. Cell 30, 343–352 (2014).

32. Conti, D. et al. Kinetochores attached to microtubule-ends are stabilised by Astrin bound PP1 to ensure proper chromosome segregation. Elife 8, (2019).

33. Meraldi, P., Draviam V. M. Sorger, P. K. Timing and checkpoints in the regulation of mitotic progression. Dev. Cell 7, 45–60 (2004).

34. Gutéerrez-Caballero, C., Burgess, S. G., Bayliss, R. Royle, S. J. TACC3-ch-TOG track the growing tips of microtubules independently of clathrin and Aurora-A phosphorylation. Biol. Open 4, 170–179 (2015).

35. Schmidt, J. C. et al. Aurora B kinase controls the targeting of the Astrin-SKAP complex to bioriented kinetochores. J. Cell Biol. 191, 269–280 (2010).

36. Chen, Q., Zhang, X., Jiang, Q., Clarke P. R. Zhang, C. Cyclin B1 is localized to unattached kinetochores and contributes to efficient microtubule attachment and proper chromosome alignment during mitosis. Cell Res. 18, 268–280 (2008).

37. Cimini, D., Wan, X., Hirel C. B. Salmon E. D. Aurora kinase promotes turnover of kinetochore microtubules to reduce chromosome segregation errors. Curr. Biol. 16, 1711–1718 (2006).

38. Manning, A. L. et al. CLASP1, astrin and Kif2b form a molecular switch that regulates kinetochore-microtubule dynamics to promote mitotic progression and fidelity. EMBO J. 29, 3531–3543 (2010).

39. Bakhoum, S. F., Thompson, S. L., Manning, A. L. Compton, D. A. Genome stability is ensured by temporal control of kinetochore-microtubule dynamics. Nat. Cell Biol. 11, 27–35 (2009).

40. Kern, D. M., Monda, J. K., Su, K.-C., Wilson-Kubalek, E. M. Cheeseman, I. M. Astrin-SKAP complex reconstitution reveals its kinetochore interaction with microtubule-bound Ndc80. Elife 6, (2017).

41. Garvanska D. H. Nilsson, J. Specificity determinants of phosphoprotein phosphatases controlling kinetochore functions. Essays Biochem. 64, 325–336 (2020).

42. DeLuca K. F., Lens S. M. A. DeLuca J. G. Temporal changes in Hec1 phosphorylation control kinetochore–microtubule attachment stability during mitosis. J. Cell Sci. 124, 622–634 (2011).

43. Huis In ‘t Veld, P. J. et al. Molecular basis of outer kinetochore assembly on CENP-T. Elife 5, (2016).

44. Zhang, G., Lischetti, T. Nilsson, J. A minimal number of MELT repeats supports all the functions of KNL1 in chromosome segregation. J. Cell Sci. 127, 871–884 (2014).

45. Vleugel, M. et al. Sequential multisite phospho-regulation of KNL1-BUB3 interfaces at mitotic kinetochores. Mol. Cell 57, 824–835 (2015).

46. Zhang, G. et al. Bub1 positions Mad1 close to KNL1 MELT repeats to promote checkpoint signalling. Nat. Commun. 8, 15822 (2017).

47. Craske, B. Welburn J. P. I. Leaving no-one behind: how CENP-E facilitates chromosome alignment. Essays Biochem. 64, 313–324 (2020).

48. Janczyk, P. L . et al. Mechanism of Ska Recruitment by Ndc80 Complexes to Kinetochores. Dev. Cell 41, 438–449.e4 (2017).

49. Radford, S. J., Hoang, T. L., Gluszek A. A., Ohkura, H. McKim K. S. Lateral and End-On Kinetochore Attachments Are Coordinated to Achieve Biorientation in Drosophila Oocytes. PLoS Genet. 11, e1005605 (2015).

50. Vallot, A. et al. Tension-Induced Error Correction and Not Kinetochore Attachment Status Activates the SAC in an Aurora-B/C-Dependent Manner in Oocytes. Current Biology vol. 28 130–139.e3 (2018).

51. Corrigan, A. M. et al. Automated tracking of mitotic spindle pole positions shows that LGN is required for spindle rotation but not orientation maintenance. Cell Cycle 12, 2643–2655 (2013).

52. Iorio, F. et al. A Semi-Supervised Approach for Refining Transcriptional Signatures of Drug Response and Repositioning Predictions. PLoS One 10, e0139446 (2015).

53. Patel, H. et al. Kindlin regulates microtubule function to ensure normal mitosis. Journal of Molecular Cell Biology, Vol 8, issue 4 (2016).

54. Shrestha, R.L. et al., TAO1 kinase maintains chromosomal stability by facilitating proper congression of chromosomes. https://doi.org/10.1098/rsob.130108 Open biology, (2014).

55. Zulkipli et al., Spindle rotation in human cells is reliant on a MARK2-mediated equatorial spindle centering mechanism. Journal of Cell Biology 217 (9), 3057–3070.

